# YprA family helicases provide the missing link between diverse prokaryotic immune systems

**DOI:** 10.1101/2025.09.15.676423

**Authors:** Ryan T. Bell, Thomas Gaudin, Yi Wu, Sofya K. Garushyants, Harutyun Sahakyan, Kira S. Makarova, Yuri I. Wolf, Simon A. Jackson, Franklin L. Nobrega, Chase L. Beisel, Eugene V. Koonin

**Affiliations:** Division of Intramural Research, National Library of Medicine, National Institutes of Health, Bethesda, United States; Helmholtz Institute for RNA-based Infection Research (HIRI), Helmholtz Centre for Infection Research (HZI), 97080 Würzburg, Germany; School of Biological Sciences, University of Southampton, SO17 1BJ Southampton, UK; School of Pharmacy and Biomedical Sciences, Division of Health, University of Waikato, Hamilton 3126, New Zealand; Medical faculty, University of Würzburg, 97080 Würzburg, Germany

## Abstract

Bacteria and archaea possess an enormous variety of antivirus immune systems that often share homologous proteins and domains, some of which contribute to diverse defense strategies. YprA family helicases are central to widespread defense systems DISARM, Dpd, and Druantia. Here, through comprehensive phylogenetic and structural prediction analysis of the YprA family, we identify several major, previously unrecognized clades, with unique signatures of domain architecture and associations with other genes. Each YprA family clade defines a distinct class of defense systems, which we denote ARMADA (dis<u>ARM</u>-related <u>A</u>ntiviral <u>D</u>efense <u>A</u>rray), BRIGADE (<u>B</u>ase hypermodification and <u>R</u>estriction Involving <u>G</u>enes encoding <u>A</u>RMADA-like and <u>D</u>pd-like <u>E</u>ffectors), or TALON (<u>T</u>OTE-like and <u>A</u>RMADA-<u>L</u>ike <u>O</u>peron with <u>N</u>uclease). In addition to the YprA-like helicase, ARMADA systems share two more proteins with DISARM. However, ARMADA YprA homologs are most similar to those of Druantia, suggesting ARMADA is a ‘missing link’ connecting DISARM and Druantia. We show experimentally that ARMADA protects bacteria against a broad range of phages via a direct, non-abortive mechanism. We also discovered multiple families of satellite phage-like mobile genetic elements that often carry both ARMADA and Druantia Type III systems and show that these can provide synergistic resistance against diverse phages.

## Introduction

Genetic parasites, and in particular, viruses are associated with all life forms and exhibit enormous diversity. The incessant host-parasite arms race, a key aspect of evolution over the past 4 billion years, has driven the emergence of complex and diverse defensive strategies, with numerous, intricate interconnections through gene and domain shuffling, complex formation, pattern recognition, synergy, signaling, co-regulation, and especially in prokaryotes, horizontal gene transfer (HGT) (Makarova, Anantharaman et al. 2014, Koonin, Makarova et al. 2017, Koonin and Krupovic 2019, Bell, Wolf et al. 2020, Bernheim and Sorek 2020, Gao, Altae-Tran et al. 2020, Makarova, Timinskas et al. 2020, Makarova, Wolf et al. 2020, Botelho 2023, Beamud, Benz et al. 2024, Bell, Sahakyan et al. 2024, Hossain, Aslan et al. 2024, Wu, Garushyants et al. 2024). The discovery of defense islands—regions of prokaryotic genomes that are significantly enriched in antiviral defense systems (Makarova, Wolf et al. 2011)— facilitated systematic identification of many distinct defense systems via so-called ‘guilt-by-association’ approaches (Doron, Melamed et al. 2018, Gao, Altae-Tran et al. 2020). However, other approaches to identifying diverse defense systems demonstrate that our understanding of the immune repertoire of prokaryotes is far from complete (Vassallo, Doering et al. 2022, Bell, Sahakyan et al. 2024).

Many distinct types of defense systems share homologous proteins or domains, indicative of common evolutionary origins and a tendency for certain protein families to diversify or be co-opted into diverse immune systems. For example, DISARM (<u>D</u>efense Island <u>S</u>ystem <u>A</u>ssociated with <u>R</u>estriction-<u>M</u>odification), Dpd (7-<u>d</u>eaza<u>p</u>urine in <u>D</u>NA), and Druantia, which provide broad protection against bacteriophages (Thiaville, Kellner et al. 2016, Doron, Melamed et al. 2018, Ofir, Melamed et al. 2018), share a gene encoding a helicase of the YprA-like family (denoted DrmAB, DpdJ, and DruE, respectively). In our recent study of Type IV restriction McrBC and CoCoNuT (<u>co</u>iled-<u>co</u>il <u>nu</u>clease tandem) systems, we detected a DISARM-like gene cassette, encoding a YprA-like helicase, frequently associated with canonical, non-CoCoNuT McrBC operons (Bell, Sahakyan et al. 2024). This observation pointed to uncharted diversity remaining in the YprA-containing defense landscape.

YprA-like helicases belong to the helicase superfamily 2 (SF2), and in addition to the canonical RecA1 and RecA2 domains, contain a characteristic zinc-binding C-terminal domain, known variously as DUF1998, MZB (<u>M</u>rfA <u>Z</u>n^2+^-<u>b</u>inding), and HAR (<u>H</u>elicase <u>A</u>llosteric <u>R</u>elay) (Gorbalenya and Koonin 1993, Fairman-Williams, Guenther et al. 2010, Roske, Liu et al. 2021, Bravo, Aparicio-Maldonado et al. 2022). The earliest biochemical characterization of a YprA homolog, to our knowledge, is a study of *M. smegmatis* SftH (<u>s</u>uperfamily two <u>h</u>elicase), which is a monomeric 3⍰ to 5⍰ helicase acting on single-stranded DNA (Yakovleva and Shuman 2012). YprA family helicases were implicated in DNA repair through a forward genetic screen in Bacillus subtilis, which identified the yprAB gene pair as critical for survival under mitomycin C (MMC) treatment (Burby and Simmons 2019). Further characterization of this operon, renamed mrfAB (<u>m</u>itomycin <u>r</u>epair factors A and B), determined that it encodes a 3⍰ to 5⍰ DNA helicase (MrfA) and a 3⍰ to 5⍰ DNA exonuclease (MrfB) that function in nucleotide excision repair (Bargonetti, Champeil et al. 2010, Burby and Simmons 2019, Roske, Liu et al. 2021). MrfAB homologs have a broad phyletic distribution, but their function appears to be species-specific, as complementation requires closely related alleles (Burby and Simmons 2019). Some MrfAB homologs in eukaryotes, in particular, plants and fungi, also provide protection against MMC DNA damage (Roske, Liu et al. 2021). Structural studies have revealed unique features of MrfA, and the DrmAB helicase encoded by DISARM, which support their distinct roles in nucleic acid processing and defense (Roske, Liu et al. 2021, Bravo, Aparicio-Maldonado et al. 2022). This extraordinary structural and functional plasticity, combined with the presence of YprA homologs across diverse defense systems present in many prokaryotic phyla, prompted us to undertake a comprehensive phylogenetic analysis of the YprA family.

Here, we report two major, previously uncharacterized branches of YprA-containing defense systems that we denote ARMADA (DIS<u>ARM</u>-related <u>a</u>ntiviral <u>d</u>efense <u>a</u>rray) systems. Additionally, we identify two minor, distinct branches of systems that we term BRIGADE (<u>B</u>ase hypermodification and <u>R</u>estriction Involving <u>G</u>enes encoding <u>A</u>RMADA-like and <u>D</u>pd-like <u>E</u>ffectors) and TALON (<u>T</u>OTE-like and <u>A</u>RMADA-<u>L</u>ike <u>O</u>peron with <u>N</u>uclease). We also detect unappreciated diversity among Druantia Types I and II, which we recognized as constituting a fourth group of CoCoNuT systems (Bell, Sahakyan et al. 2024). These encode inactivated homologs of the McrB-like GTPase CnuB, the active version of which is present in all previously described CoCoNuTs. We further analyze YprA homologs from each major phylogenetic branch and detect several domains that distinguish these branches and likely contribute to their functional specialization. We validate the functionality of ARMADA as an anti-phage defense system that protects bacteria against infection without causing cellular growth arrest and show that it synergizes with co-encoded Druantia Type III to expand the cell’s defensive capacity. Lastly, we find that the tested ARMADA and Druantia Type III systems are jointly embedded in defense islands that comprise cargo of distinct putative mobile genetic elements (MGEs), a characteristic of defense islands that appears to be widespread in prokaryotes.

## Results and Discussion

### New classes of defense systems encoding YprA-like proteins

We searched for YprA homologs using PSI-BLAST against the NCBI non-redundant protein sequence database and a database of completely sequenced prokaryotic genomes maintained by our group. After applying several filtering strategies (see ‘Methods’), we retrieved nearly 52,000 YprA-like proteins, distributed broadly among prokaryotes (Supplementary Table S1). These YprA-like helicases have undergone extraordinary evolutionary diversification, giving rise to numerous clades with broadly variable genomic contexts (Figure 1). There are two main branches, one comprising DNA repair-related helicases either with no conserved genomic context (“YprA only,” Figure 1, no coloration) or associated with (or fused to) MrfB/YprB homologs (Figure 1, purple and gray), and a second, highly ramified branch including helicases found in several conserved operons, some corresponding to known anti-phage defense systems, including DISARM, Dpd, and Druantia/Type IV CoCoNuT) (Figure 1, red, cyan, yellow, green, maroon, black). We hypothesized that the uncharacterized clades of the second branch also represent distinct, previously unknown defense systems. The two largest uncharacterized YprA-like clades encompass a distinct set of related operons that, in addition to the YprA homologs, share two other genes with DISARM Class I systems; both encode homologs of the BREX-like BrxHII/RapA-like helicase and PglX-like methyltransferase. Therefore, we named these predicted defense systems ARMADA (dis<u>ARM</u>-related <u>A</u>ntiviral <u>D</u>efense <u>A</u>rray) (Figure 1, magenta, blue).

**Figure 1.**
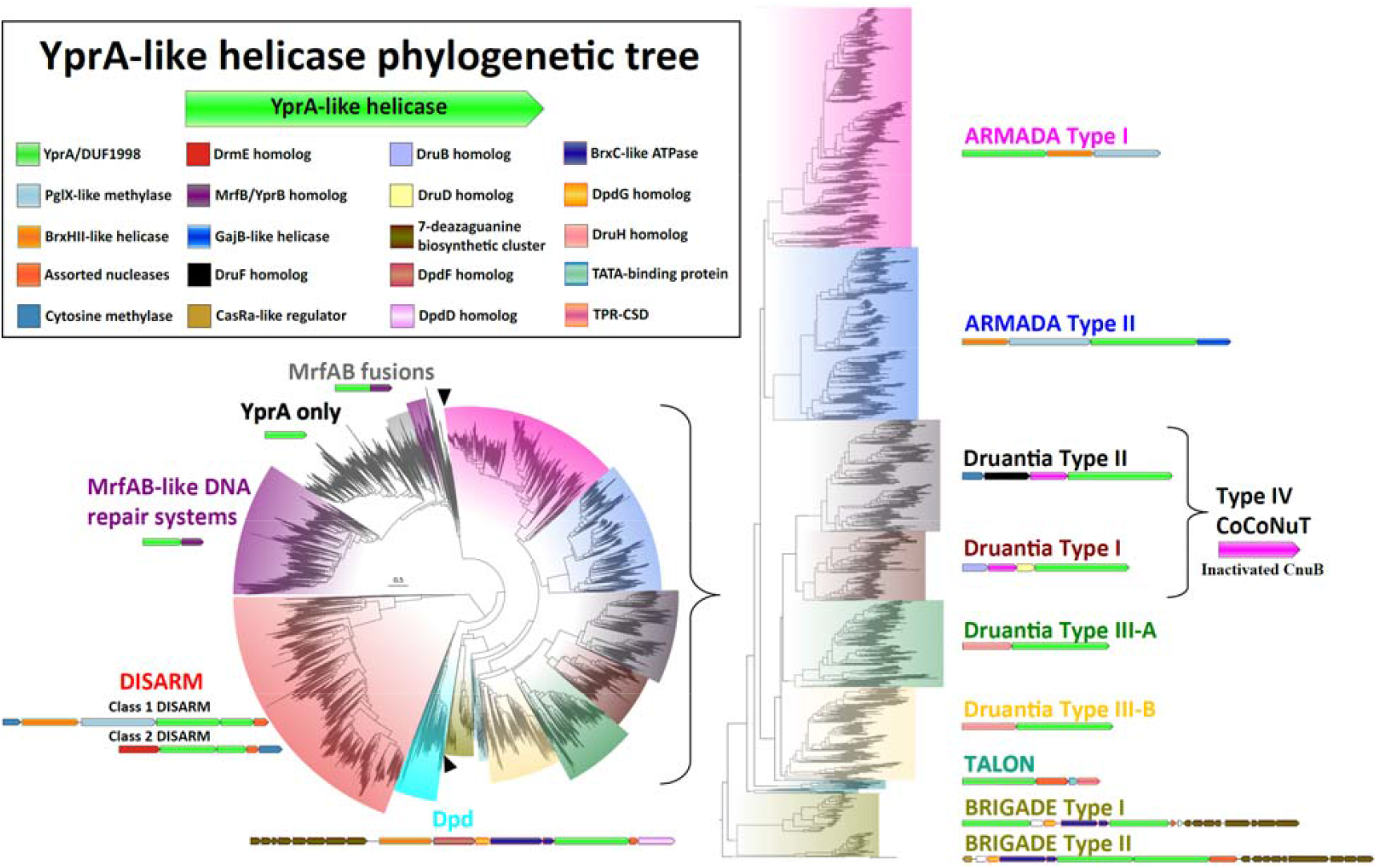
YprA-like helicase phylogeny reveals multiple, distinct defense system classes. The YprA family of helicases consists of two distinct branches, those involved in defense (red, cyan, teal, yellow, green, maroon, black, blue, and magenta), and those involved in DNA repair and perhaps other housekeeping functions (gray and purple). The DISARM-like clade, with 96% bootstrap support, encompasses the DISARM and Dpd defense systems, as well as many uncharacterized DISARM-like and Dpd-like classes. The other defensive clade encompasses the ARMADA and Druantia systems and is enlarged for clarity, with black wedges indicating the boundaries of the enlarged portion of the full tree. Each of the branches with leaves all of one color is labeled, using a font with the same color, with the most abundant defense system found in that branch, as well as a representative operon. These distinct conserved genomic associations are highly abundant within but not perfectly confined to the respective groups. This tree was built from the representatives of 90% identity clusters of all validated YprA homologs.

The ARMADA clade splits into two well-separated, strongly supported subclades which we denote Type I and Type II. Both ARMADA types are most abundant in Actinomycetota and Pseudomonadota, but Type I shows a wider phyletic distribution, including a variety of archaea, whereas Type II is absent from archaea (Supplementary Table S1). ARMADA Type I is generally arranged with a gene encoding the YprA homolog first, followed by the BrxHII-like and PglX-like methyltransferase genes. However, this order is reversed in many archaea and cyanobacteria, with the BrxHII-like helicase and PglX-like methylase gene pair preceding the YprA-like helicase, resembling ARMADA Type II. In contrast, ARMADA Type II exhibits a strict gene order and additional complexity, with the aforementioned arrangement almost always followed by a GajB-like helicase (Doron, Melamed et al. 2018) fused at its N-terminus to a RelE-like predicted RNase toxin domain (Supplementary Table S2).

In this same clade with ARMADA and Druantia, we also detected two smaller, deeper-branching lineages. One represents a group of related systems that share several homologs with Dpd systems, including a predicted 7-deazaguanine derivative biosynthetic cluster and typically Phospholipase D (PLD) family endonucleases, although occasionally these are substituted with PD-(D/E)xK family endonucleases. There appear to be two types of these Dpd-like systems, both usually encoding two YprA-like helicases. The Type I systems encode both an ARMADA-like YprA homolog and a YprA family helicase that is found in the Dpd-containing branch of the YprA tree. For the Type II systems, the two YprA homologs are nearly identical and are both ARMADA-like. We dubbed these systems BRIGADE (<u>B</u>ase hypermodification and <u>R</u>estriction Involving <u>G</u>enes encoding <u>A</u>RMADA-like and <u>D</u>pd-like <u>E</u>ffectors), as they appear to be hybrids that combine features of both Dpd and ARMADA (Supplementary Figure S1). BRIGADE systems were detected exclusively in halophilic archaea (Supplementary Table S1). Finally, we detected an extremely rare system in Bacillati, largely confined to Chloroflexota, which typically encodes an ARMADA-like YprA homolog and a PD-(D/E)xK family endonuclease of the NERD (<u>N</u>ucleas<u>e</u>-<u>R</u>elated <u>D</u>omain) subtype (Grynberg and Godzik 2004, Steczkiewicz, Muszewska et al. 2012), as well as a tetratricopeptide repeat-cold shock domain fusion (TPR-CSD) and TATA-binding proteins (TBP). These latter two genes are shared with the TOTE (<u>T</u>PR, <u>O</u>B, <u>T</u>BP, <u>E</u>ffector) family of conflict systems (Aravind, Iyer et al. 2022). We denote these as TALON (<u>T</u>OTE-like and <u>A</u>RMADA-<u>L</u>ike <u>O</u>peron with <u>N</u>uclease) systems. Collectively, the ARMADA, BRIGADE, and TALON systems account for the majority of the as-yet unclassified YprA-like homologs in the ‘defense’ branch of the YprA tree.

### Anti-phage activity of ARMADA

To experimentally validate ARMADA as an anti-phage defense system, we selected an ARMADA Type II cluster from *E. coli* NCTC 12900 (Figure 2A). This gene cluster encodes four proteins, namely, three predicted helicases (ArmA, ArmC, ArmD) and one predicted methyltransferase (ArmB). Specifically, ArmA is a BrxHII-like helicase with a DEXD RapA (Swi2/Snf2) domain; ArmB is a PglX-like DNA methyltransferase (YeeA-related); ArmC is a YprA-like ATP-dependent DEAD/DEAH-box helicase; and ArmD is a GajB-like UvrD family helicase with an N-terminal RelE-like RNase domain (Supplementary Table S2). We cloned the full cluster under its native promoters into plasmid pBeloBAC11 and assessed anti-phage activity against a panel of 80 diverse phages (Supplementary Tables S3-5). ARMADA Type II conferred protection (defined as >10^1^ PFU/ml reduction) against 13 of these phages, with the strongest activity observed against phage HK022 (Figure 2B, Supplementary Figure S3, Supplementary Table S6).

**Figure 2.**
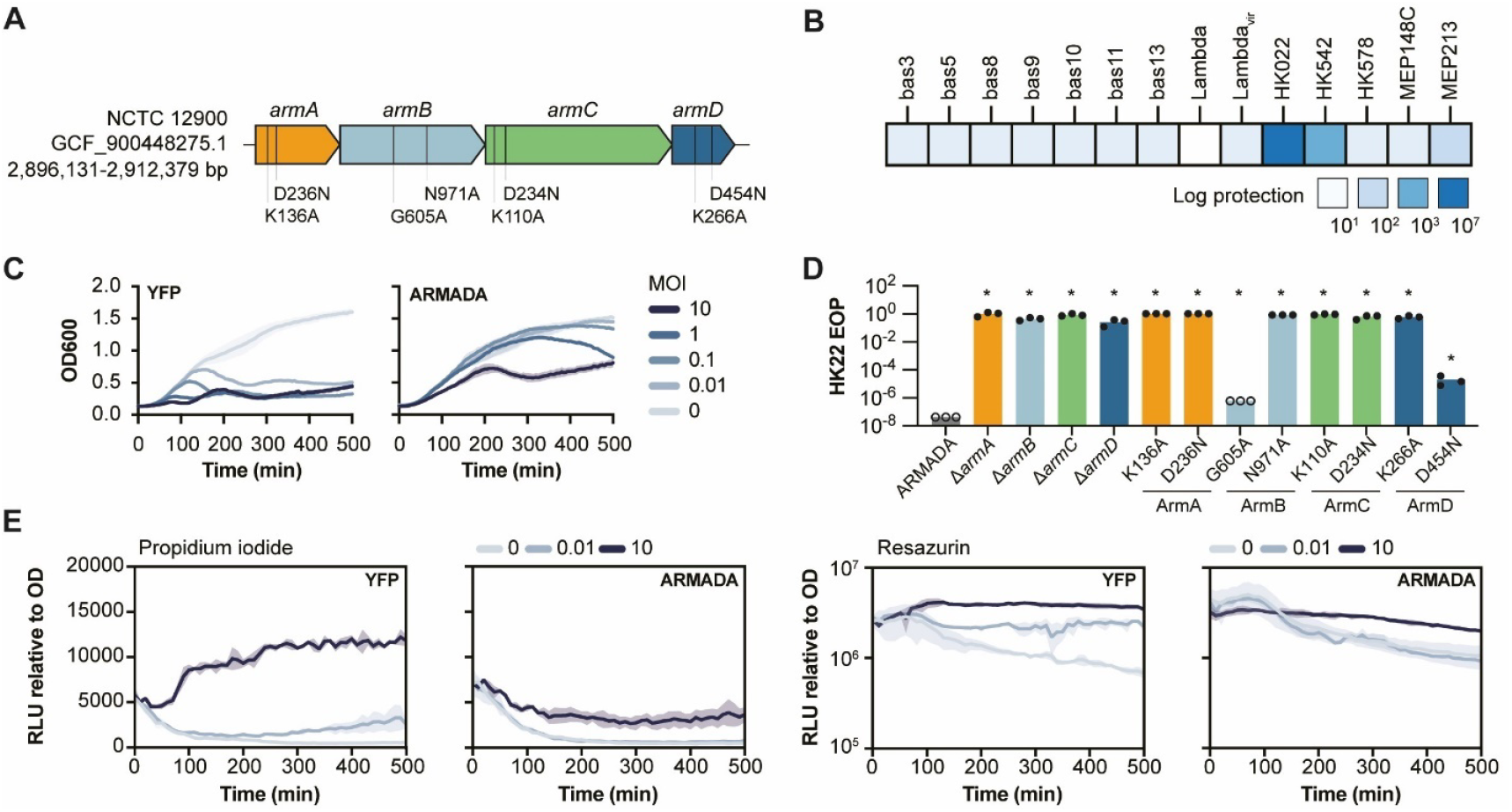
Experimental validation of anti-phage activity of ARMADA. (A) Schematic of the ARMADA Type II cluster from *E. coli* NCTC 12900 (GCF_900448275.1), showing gene architecture and mutated residues used in (D). Genes are colored as in Figure 1. (B) Log-protection (reduction in phage titer) conferred by ARMADA, expressed from plasmid pBeloBAC11 in BL21-AI, against a panel of 80 phages (Supplementary Figure S3, Supplementary Table S6). A representative subset against which ARMADA provides protection is shown. (C) Growth curves of *E. coli* expressing yellow fluorescent protein (YFP) control (left) or ARMADA (right) upon infection with phage HK022 at increasing multiplicity of infection (MOI), measured as optical density (OD_600_) over time. Curves show mean and standard deviation from three independent experiments. (D) Efficiency of plating (EOP) of phage HK022 on cells expressing wild-type ARMADA, individual gene deletions, or point mutants. Data represent three independent experiments. * Statistically significant differences to ARMADA control, p < 0.05 (see Methods). (E) Membrane integrity (cell viability) and metabolic activity of control (YFP) and ARMADA cells infected with phage HK002. Propidium iodide fluorescence (Relative light units (RLU)/OD_600_, left) reports membrane permeabilization and cell death. Resazurin fluorescence (Relative light units (RLU)/OD_600_, right) reports redox metabolic activity. Curves show mean and standard deviation from three independent experiments.

In liquid culture, ARMADA-expressing cells showed robust protection against HK022 at both high and low multiplicity of infection (MOI), although cell lysis was still apparent at the highest MOIs (Figure 2C). Deletion of any of the four genes comprising this system abolished protection, indicating that all four components are essential for defense (Figure 2D). Mutational analysis of conserved amino acids further supported the functional importance of the enzymatic domains of the four proteins. Disrupting Walker A or B motifs in ArmA (K136A, D236N) and ArmC (K110A, D234N), Walker A motif in ArmD (K266A), and the NPPW methyltransferase motif in ArmB (N971A) abolished protection. Mutation of the Walker B motif in ArmD (D454N) and the S-adenosylmethionine (SAM)-binding GxG motif (G605A) in ArmB resulted in only partial loss of activity, suggesting these motifs contribute to, but are not essential for, the defense function of ARMADA (Figure 2D).

To further characterize ARMADA-mediated protection, we performed propidium iodide (PI) and resazurin assays to monitor cell viability and cellular metabolism during infection (Figure 2E). PI is a membrane-impermeant dye that fluoresces upon binding DNA, serving as a marker of membrane permeabilization and cell death. In control cells, high-MOI infection triggered a strong PI signal, consistent with phage-induced lysis. In contrast, ARMADA-expressing cells showed minimal PI uptake, indicating that the cells remained viable and intact following infection. Resazurin is a redox-sensitive dye that reports metabolic activity through fluorescence. In control cells, HK022 infection triggered a strong increase in resazurin signal, suggestive of phage-induced metabolic hijacking. This increase was largely absent in ARMADA-expressing cells, suggesting that ARMADA blocks phage replication without triggering shutdown or reprogramming of the host metabolism.

### Structure predictions and domain architectures of YprA homologs

To examine how YprA-like homologs diversified within distinct defense systems, such as ARMADA, we next investigated the domain architectures of the respective YprA homologs. Previous comparison of the YprA family to the SF2 helicases revealed several YprA-specific features, including an oligonucleotide/oligosaccharide-binding (OB) fold and an antiparallel beta-sheet connector element (CON) (Roske, Liu et al. 2021) that links the OB fold to the zinc-binding DUF1998 domain. Typically, the OB-CON-DUF1998 region is fused to the helicase core, but in DISARM systems it is encoded by a separate, adjacent gene (Ofir, Melamed et al. 2018).

Structural analyses of DISARM confirmed complex formation between the YprA-like helicase DrmA and the OB-CON-DUF1998-containing protein DrmB (Supplementary Figure S2) (Bravo, Aparicio-Maldonado et al. 2022). In the YprA family helicase MrfA, the OB-CON-DUF1998 multidomain module forms a canopy over the RecA-like domains and undergoes a prominent conformational change upon ssDNA binding (Supplementary Figure S2A) (Roske, Liu et al. 2021). As the ATP hydrolysis-powered helicase core inchworms along a DNA strand, the shift in the canopy position periodically entraps the substrate, in sync with the corresponding interpolations of the RecA2 domain, by tautening an interdomain linker that is predicted to ratchet through individual nucleobases, likely contributing to DNA translocation (Roske, Liu et al. 2021). Superposition of the DrmA structure onto MrfA (Supplementary Figure S2B-D) reveals the OB fold region as having diverged substantially between MrfA and DrmB, whereas the rest of the CON/DUF1998 structure appears to be conserved (Supplementary Figure S2B-D). These observations suggest that the translocation mechanism of DrmA is similar to that of MrfA, whereas the structure of the targeted DNA, likely bound by the OB fold, is distinct.

With these experimental structures as guides, we predicted structures of representatives of each major YprA clade and demarcated their globular domains via structure-based searches for matches to characterized folds (see Methods) (Figure 3, Supplementary Table S2). Beginning at the N-terminus, all YprA homologs contain a beta-strand appended at or near the N-terminus, preceding RecA1 but structurally inserted into the beta-sheet of RecA2; this element was previously observed in MrfA and denoted NTR (<u>N</u>-terminal <u>r</u>egion) (Figure 3) (Roske, Liu et al. 2021). Immediately after RecA1, YprA homologs from ARMADA, BRIGADE, TALON, and Druantia contain an inserted helical domain resembling BrxA (a small, DNA-binding protein found in several BREX antiviral system types) (Goldfarb, Sberro et al. 2015) (Figure 3 in yellow, Supplementary Table S2). This domain more closely resembles BrxA in the deeper branching ARMADA and BRIGADE systems than that of the later-branching Druantia systems (Figure 3, Supplementary Table S2). It appears likely that this BrxA-like domain originated from a single insertion into a YprA family helicase that became the founding member of the ARMADA-BRIGADE-TALON-Druantia lineage and thus represents a synapomorphy of this lineage, supporting its monophyly.

**Figure 3.**
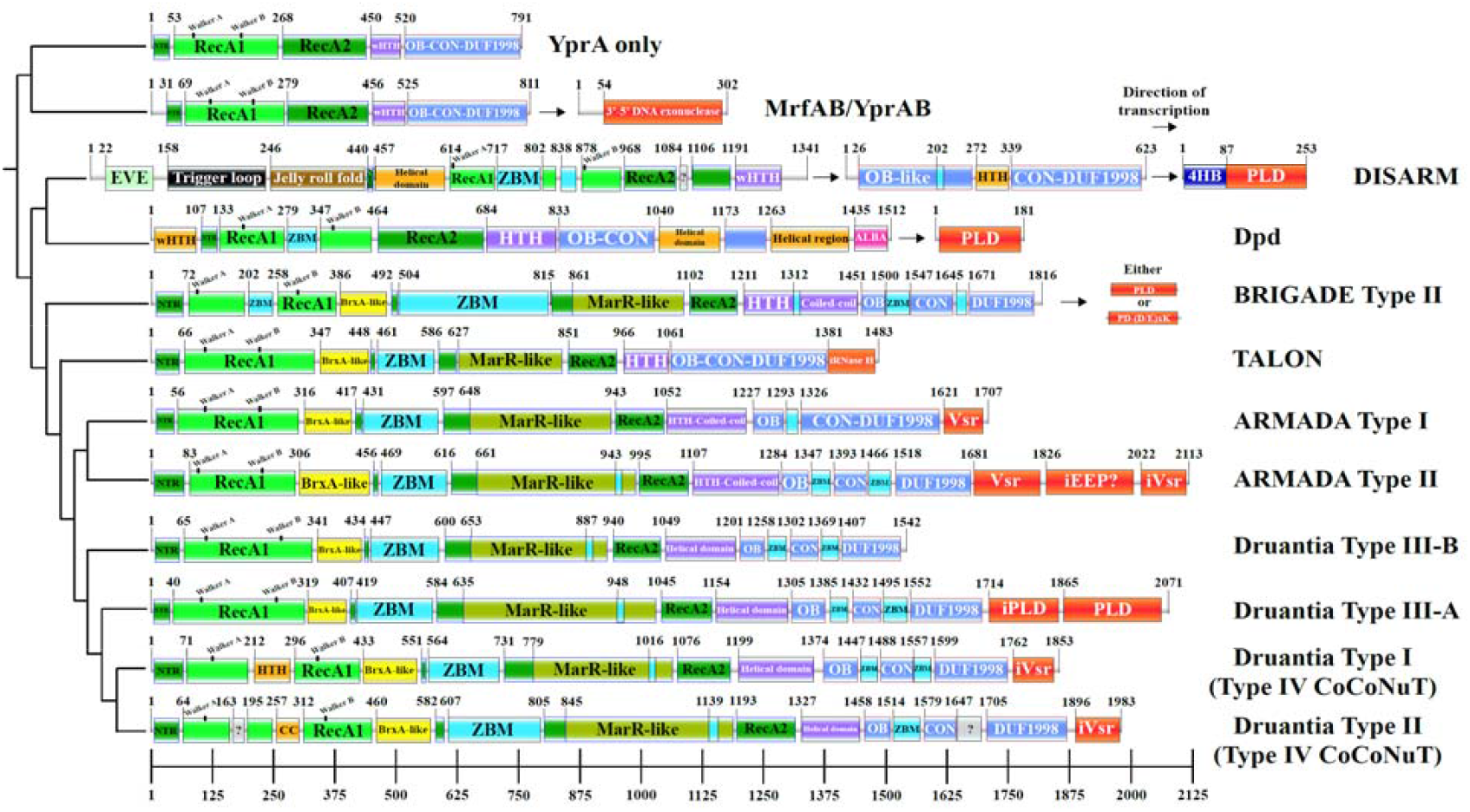
Representative domain architectures of YprA family helicases and association with nucleases. Using AlphaFold3, structures of several representative YprA family helicases from each major branch were predicted and the high-quality models (average predicted local distance difference test [pLDDT] > 80) were superimposed to select the best representative for each branch. The boundaries of globular domains were determined by comparing amino acid sequences to each structural model, as well as using searches of subsets of the models against the PDB using DALI to zero in on regions that harbored distinct folds. The sequences were also analyzed in detail using HHpred, which revealed many zinc-binding motifs. The sequences used for each representative are: UER53415.1 (YprA only), MBL8484594.1, MBL8484593.1 (MrfAB/YprAB), EHM9959445.1, EHM9959446.1, EHM9959447.1 (DISARM), EGG1131020.1 (Dpd), WP_226013624.1 (BRIGADE), GHO49638.1 (TALON), WP_108720193.1 (ARMADA Type I), WP_000937974.1 (ARMADA Type II), GIH50614.1 (Druantia Type III-B), MES1925543.1 (Druantia Type III-A), WP_061581103.1 (Druantia Type I), KQQ96139.1 (Druantia Type II). The tree is based only on the alignment of YprA-like helicases, but in some cases where the helicase is split into multiple genes, these separate gene products are shown in their operonic order to facilitate comparison of the compositions of different systems.

Consistent with having a distinct common ancestor, the domain architectures of the YprA-like helicases of ARMADA, BRIGADE, TALON, and Druantia are similar from the BrxA-like domain through the end of RecA2, especially when compared to the DISARM and Dpd YprA-like proteins. All these apparently more recently diverged homologs contain a predicted beta-barrel domain inserted into the start of the RecA2 domains, adjacent to the BrxA-like domain (Figure 3 in cyan, Supplementary Table S2). This beta-barrel domain contains at least two, and often more, conserved CxxC predicted Zn^2+^-binding motifs, some of which resemble RanBP2-type zinc fingers, whereas others are similar to TerY-C metal binding domains (Supplementary Table S2). It is one of several domains in the various defensive YprA-like helicases that have evolved conserved motifs of this type (see below), which we collectively denote <u>z</u>inc-<u>b</u>inding <u>m</u>odules (ZBMs). Adjacent to the beta-barrel ZBM and also inserted into the RecA2 fold is a MarR-like domain. These transcriptional regulator-like domains usually harbor additional ZBMs near their C-termini (Figure 3 in olive, cyan, Supplementary Table S2). The YprA homologs in BRIGADE Type II and ARMADA Types I and II also contain HTH-like domains with large, finger-like helical extensions that are predicted to be coiled-coils, between the RecA2 fold and the start of the OB-CON-DUF1998 domain array (Figure 3 in purple). These helical extensions occupy the same position, directly following the RecA2 fold, as a variety of other (typically, much smaller) helical domains found in all other YprA homologs, suggesting they all are homologous despite low sequence similarity (Figure 3 in purple). This region in BRIGADE Type II also contains a ZBM (Figure 3 in cyan).

The canopy-forming OB-CON-DUF1998 modules (see above) in the defensive YprA-like helicases, compared to their homologs in the YprA-only and MrfAB/YprAB branches, are interrupted by small domains, particularly in the OB fold region, which almost always harbor additional predicted ZBMs (Figure 3 in blue and cyan). There are two of these inserted domains in all Druantia types, ARMADA Type II, and BRIGADE Type II, and one in ARMADA Type I (Figure 3). In Dpd, this region contains two larger helical insertions, as well as an ALBA-like (<u>A</u>cetylation lowers <u>b</u>inding <u>a</u>ffinity) nucleic acid-binding domain fused at the C-terminus (Figure 3, Supplementary Table S2) (Goyal, Banerjee et al. 2016). Although the defensive YprA homologs as a whole are characterized by a dramatic increase in the content of predicted ZBMs, we detected only one in canonical Dpd systems, inserted into the first RecA-like fold. Furthermore, we found that the 4-cysteine zinc-binding cluster (DUF1998), which characterizes the entire YprA family, was lost entirely in Dpd (although some deeper branching Dpd-like systems retain it). In DISARM, where the canopy-forming module is encoded in a separate gene, it is interrupted by a HTH domain-containing, predicted DNA-binding domain between the OB fold and the CON beta-sheet, and also contains a ZBM (Figure 3, Supplementary Table S2). These distinct, complex elaborations, often containing ZBMs, might have evolved in defensive YprA-like helicases as part of the arms race, to keep up with susceptible phages. This architectural plasticity might be one of factors underlying the selection of this particular helicase family for central roles in diverse antiviral defense systems. Where present, the ZBMs likely aid in target recognition, stabilization of the large helicase structure, and/or complex formation with other components of the defense systems.

### Nucleases associated with YprA family helicases

An important distinction between the main branches of YprA homologs is the presence of a variety of active and inactivated nuclease domains, either fused to the helicases at their C-termini or encoded by downstream genes in the respective operons (Figure 3, in red). As described above, MrfB in the MrfAB systems contains a 3’ to 5’ exonuclease domain. DISARM systems typically include a PLD superfamily endonuclease encoded following the DUF1998-containing protein (DrmC); most of these nucleases contain a 4-<u>h</u>elix <u>b</u>undle (4HB) at the N-terminus with structural similarity to Cas11 proteins (Figure 3, Supplementary Table S2). The DISARM systems encompass the only branch of YprA homologs where the helicase and the OB-CON-DUF1998 domain array are consistently encoded by separate genes (although this separation is sporadically observed in other lineages, too), and the presence of the 4HB domain might be linked to this distinct operon architecture. The DISARM-like YprA family helicases in Dpd, which are not broken into two genes, are associated with a PLD-type nuclease that lacks the 4HB domain (Figure 3). BRIGADE Type I and Type II systems, which are most closely related to Dpd, also typically include either a PLD or PD-(D/E)xK-type nuclease in the neighborhood, with no 4HB domain (Figure 3, Supplementary Figure S1, Supplementary Table S2).

In contrast, the YprA family helicases of ARMADA, TALON, and Druantia typically contain a nuclease domain fused to their C-termini. Uniquely among YprA homologs, those encoded by TALON systems are fused at the C-terminus to inactivated RNase H fold domains (Supplementary Table S2), consistent with their similarity to TOTE systems, which often also include RNase H domains (Aravind, Iyer et al. 2022). In ARMADA Type I and II, the C-terminal domain is a predicted active Vsr (<u>V</u>ery <u>s</u>hort patch <u>r</u>epair)-type nuclease, which is a member of the PD-(D/E)xK superfamily (Figure 3, Supplementary Table S2) (Tsutakawa, Jingami et al. 1999). YprA homologs from ARMADA Type II often, but not invariably, contain additional predicted inactivated nuclease domains of the EEP (endonuclease/exonuclease/phosphatase) and Vsr families following the predicted active domain (Figure 3, Supplementary Table S2) (Chaudhari, Ware et al. 2021). Druantia Type III encompasses two PLD-like nuclease domains, with only the most C-terminal domain predicted to be active (Figure 3, Supplementary Table S2). YprA family helicases in the phylogenetically distinct branch of Druantia Type III-like systems (Figure 2, yellow branch) occasionally contain these domains, but in other cases, contain a Vsr nuclease or, most frequently, no nuclease domain at all (Figure 3, Supplementary Table S2). Finally, YprA homologs of Druantia Type I and II (Type IV CoCoNuT) both contain predicted inactivated Vsr C-terminal nuclease domains (Figure 3, Supplementary Table S2).

### Additional domains in YprA family helicases

We additionally detected several domains unique to various YprA-like branches. Druantia Type I and II (Type IV CoCoNuT) contain insertions of either an HTH, or two small uncharacterized domains, into their RecA1 domains, respectively (Figure 3). The second insertion into the RecA1 domain in Druantia Type II is predicted to be a coiled-coil, and might play a role in interacting with the CoCoNuT CnuB homolog also encoded in this operon and containing an extended coiled-coil region (Bell, Sahakyan et al. 2024). BRIGADE Type II has a beta-hairpin ZBM inserted into the RecA1 domain (Figure 3). Dpd also contains a ZBM inserted into RecA1, as does DISARM, in which two ZBMs are embedded in the RecA1 fold, potentially revealing an underlying functional similarity between these deep-branching, otherwise radically diverged YprA homologs. YprA-like helicases of the Dpd systems encompass an additional unique domain, an N-terminal wHTH domain resembling the wHTH domain of the bacterial chromosome partitioning protein MukF (Figure 3, Supplementary Table S2).

DISARM Class 1 YprA-like helicases are the only other group of YprA homologs, besides those in Dpd systems, which contain a predicted DNA-binding domain at their N-termini, namely, an EVE-like domain that can be predicted to recognize modified bases (Figure 3, Supplementary Table S2) (Bertonati, Punta et al. 2009, Bell, Wolf et al. 2020). Following this EVE-like domain, DISARM Class 1 YprA homologs encompass a largely unstructured region that has been described as an autoinhibitory “trigger loop” partially occluding the DNA-binding site (Bravo, Aparicio-Maldonado et al. 2022). Adjacent to the trigger loop, these helicases contain a jelly roll domain which is followed by the NTR, here not located at the N-terminus, but still inserting into the RecA2 domain, and then, by a helical domain (Figure 3, Supplementary Table S2). At their C-termini, DISARM Class 1 YprA homologs usually contain another helical domain that is analogous to the wHTH/HTH domains found between RecA2 and the OB-CON-DUF1998 multidomain module in the YprA only and MrfAB/YprAB branches. As noted above, all helical domains occupying this position in the various YprA-like lineages, including those in Dpd, BRIGADE, TALON, ARMADA, and Druantia, might be distant homologs (Figure 3 in purple).

### Co-occurring ARMADA Type II and Druantia Type III form a potent synergistic anti-phage defense

We identified a genomic region in many *E. coli* isolates where three defense systems, Zorya Type II, Druantia Type III, and ARMADA Type II form a conserved array. Previous work showed that Zorya Type II and Druantia Type III systems found in one of these defense gene clusters, in *E. coli* strain ECOR19, provide synergistic protection against specific phages (Wu, Garushyants et al. 2024). We hypothesized that the entire tripartite Zorya-Druantia-ARMADA array functions in concert to enhance defense. For the initial test of this hypothesis, we explored the relationship between Druantia Type III and ARMADA Type II, choosing this pair of systems because all their genes are co-directional and they both encode YprA-like helicases.

We investigated the activities of ARMADA Type II and Druantia Type III in *E. coli* strain ATCC 8739, which encompasses the Zorya-Druantia-ARMADA locus (Figure 4A). To this end, we generated precise deletions of the respective systems, producing single and double mutants (Figure 4A, Supplementary Figure S2) and measured phage sensitivity based on efficiency of plating (EOP) and the size fold change (SFC) of the plaques using the double-deletion strain as the reference (Figure 4B). Screening 66 phages that could form plaques on the double-deletion strain showed that the strain carrying only Druantia III exhibited at least modest protection (<0.01 EOP or <0.5 SFC) against 3 phages, whereas the strain carrying ARMADA Type II exhibited a much broader range of immunity, with at least modest protection observed against 30 phages. The susceptible phages were mostly from the families Drexlerviridae and Demerecviridae. Notably, the wild-type (WT) *E. coli* strain ATCC 8739, carrying both Druantia III and ARMADA Type II, exhibited enhanced protection against multiple tested phages compared to the deletion strains carrying only a single system, suggesting potential synergistic interaction between the two systems.

**Figure 4.**
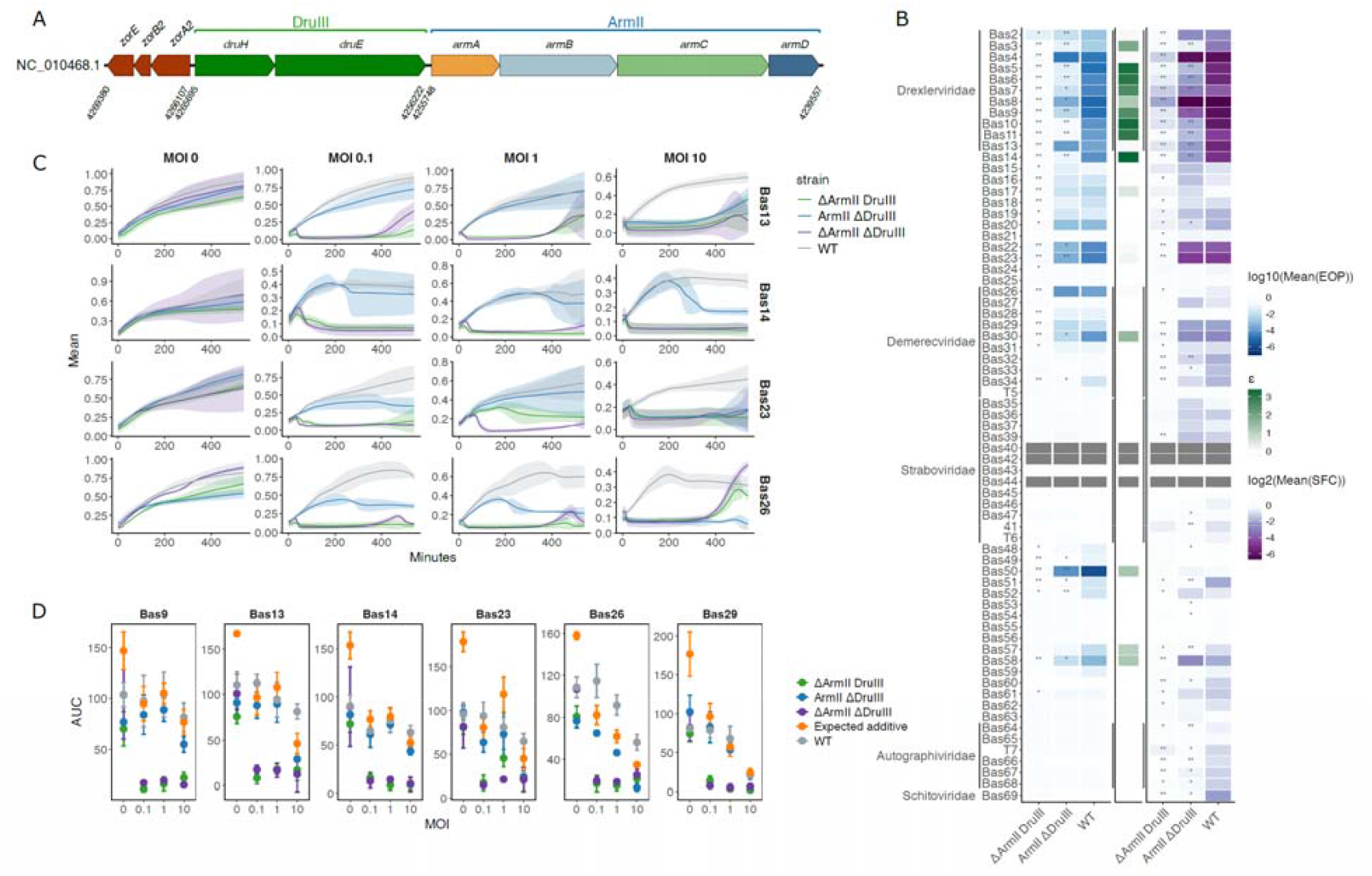
Co-occurring ARMADA Type II and Druantia Type III synergize to confer high anti-phage resistance in the native host. (A) Genetic locus encoding ARMADA Type II, Druantia Type III, and Zorya Type II in *E. coli* strain ATCC 8739 (NC_010468.1). (B) Efficiency of plating (EOP) and plaque size fold change (SFC) of phages on *E. coli* strain ATCC 8739 with the ARMADA Type II and Druantia Type III systems intact or deleted. EOP and SFC are calculated relative to the double-deletion. The epistatic coefficients (ε) are shown on the right and are calculated based on EOP (see Methods) for all systems showing at least moderate protection (< 0.01). Dark gray bars: no plaques. Heat map values represent the mean of three independent experiments. Asterisks indicate statistically significant differences in EOP or SFC compared with WT (one-sided t-test): ** p < 0.01, * p < 0.05. (C) Growth curves for 4 representative phages at different MOI. Each curve is a mean of three independent replicates, and the semi-transparent areas around the curves represent 95% confidence intervals (D) Area under growth curve (AUC) at different MOI. Expected additive (orange) is the sum of AUCs for mutants carrying only one defense system. In cases of synergy, the AUC of the WT strain (gray) is expected to be larger than expected additive.

To quantify the synergy between ARMADA and Druantia, we calculated synergy scores (epistatic coefficients (ε), see Methods) for the efficiency of plating for 21 phages where the WT strain showed at least moderately reduced plaque formation compared to the double-deletion strain (<0.01). We observed potential synergistic interactions (positive ε) for 16 phages, especially from Drexlerviridae (Figure 4B). However, in cases where no changes in plaque formation were observed, particularly Autographiviridae, Schitoviridae, and some unclassified phages, we detected significant differences in plaque sizes between individual mutants and WT, suggesting a broader interplay between the two systems (Figure 4B). Thus, Druantia Type III and ARMADA Type II systems showed clear evidence of synergizing to confer phage defense in their native host, but only for a subset of phages.

We separately explored the behavior of the deletion mutants in liquid culture using six representative phages (Bas9, Bas13, Bas14, Bas23, Bas26, Bas29) tested at different MOIs (Figure 4C and 4D). For these phages, ARMADA consistently conferred greater protection than Druantia. In the prior EOP assay, we observed high ε between ARMADA and Druantia for Bas9 and Bas14, meaning strong evidence for synergistic interaction, and small positive ε values for Bas13 and Bas23, but found no evidence of synergy for Bas26 or Bas29 (Figure 4B). In contrast, in liquid culture, the WT strain carrying both Druantia and ARMADA grew better than either of the deletion mutants in the cases of Bas13, Bas14, Bas23 and Bas26 with least one MOI, implying synergy (Figure 4C). To differentiate between simply additive and genuinely synergistic interactions, we compared the areas under the growth curve (AUC) for the combined and individual systems. Synergy was defined as occurring when the AUC of a strain carrying both systems exceeded the sum of the AUCs for the corresponding individual systems. Although this definition is deliberately stringent, we identified such cases for four phages (Bas13, Bas14, Bas23 and Bas26) (Figure 4D). For three of the phages, these observations correlated with the EOP assay results, but with a different effect magnitude, whereas Bas26 exhibited strong synergy in liquid culture that was not detected using EOP. This observation shows that detection of synergy can depend not only on the phage, but also on the types of measurements and infection conditions.

These findings underscore a critical role for the ARMADA Type II system in phage defense in *E. coli* strain ATCC 8739, demonstrating that it overlaps with, but also extends considerably beyond, the protection range of Druantia Type III. Moreover, the observed synergistic effects between the Druantia Type III and ARMADA Type II systems against certain phages suggest a complex relationship between these two systems that warrants deeper mechanistic exploration.

### Many defense islands are derived from uncharacterized integrated MGEs

During our investigations of the CoCoNuTs and now the ARMADA systems, it has become clear that most large, conserved defense islands containing these systems are associated with MGE-related genes and in many cases reside within MGEs. These MGEs are likely involved in horizontal transfer of the respective defense systems to other prokaryotic genomes, typically, utilizing a site-specific integrase and conserved integration sites found in tRNA or tmRNA genes.

In our characterization of ARMADA Type II, we tested systems from two *E. coli* strains, NCTC 12900 and ATCC 8739, each located in a Zorya-Druantia-ARMADA defense island. These two islands, despite a high degree of similarity and residing in closely related strains, have clearly distinct genomic contexts that imply they have been delivered by separate MGE types which target different tRNA genes (Figure 5A and 5B). Conservation and co-mobilization of the Zorya-Druantia-ARMADA gene cluster across these disparate neighborhoods provide further evidence of coordinated function, reinforcing the experimental demonstration of synergy between ARMADA and Druantia.

**Figure 5.**
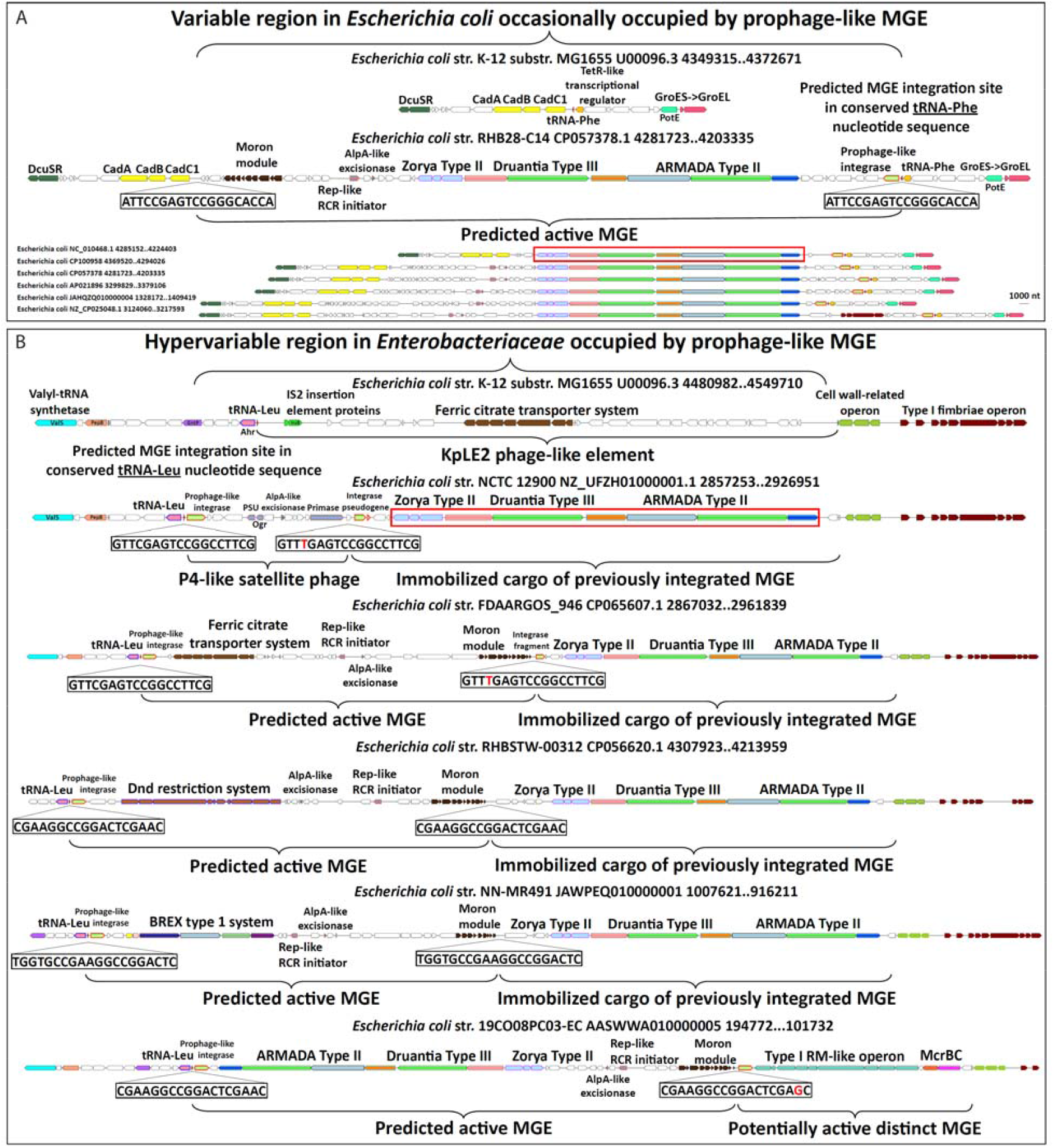
Putative MGEs carrying as cargo a conserved array of ARMADA Type II, Druantia Type III, and Zorya Type II inserted at tRNA-Phe and tRNA-Leu genes in *E. coli*. (A) Comparative genomic analysis of ARMADA and Druantia systems revealed these defense islands in *E. coli* appear to reside in large MGEs that have integrated into a tRNA-Phe gene using a prophage-like mechanism. The strain tested in Figure 4, ATCC 8739, is outlined by a red box, and is not enlarged because the MGE appears to have been immobilized. The boundaries of the integrated element are direct repeats, one in the tRNA-Phe gene at the 3’ end of the region (adjacent to an orange colored TetR-like gene) and one at the 5’ end, next to a yellow colored cadaverine synthesis operon. Additional conserved factors encoded in the flanking regions such as GroEL and DcuSR are shown as visual aids. (B) Additional ARMADA and Druantia-containing defense islands in *E. coli* that appear to be composites of large MGEs that have integrated into a tRNA-Leu gene using a prophage-like mechanism, with the ARMADA-containing island frequently, but not always immobilized. When the ARMADA-containing element is immobilized, the predicted active elements, which have likely integrated at the tRNA-Leu gene subsequently, carry other defense genes, or in the case of the strain we tested in Figure 2, NCTC 12900 (outlined by a red box), resemble a P4 satellite phage. The boundaries of the active elements are direct repeats that may have 1 mismatch (in red), one in the tRNA-Leu gene at the 5’ end of the region (adjacent to a pink colored Ahr aldehyde reductase gene) and one at the 3’ end which abuts the immobilized ARMADA-containing element. In these immobile elements, vestiges of the original delivering recombinase can often be found in the form of integrase gene fragments or pseudogenes. The entire hypervariable defense island region is bounded at the 3’ end by an operon of cell wall-related factors (green). Additional conserved factors encoded in the flanking regions such as ValS tRNA synthetase (cyan) and a Type I fimbriae operon (maroon) are shown to aid in visualizing the borders. Within the active elements are AlpA-like excisionases that facilitate recombination and Rep-like rolling circle replication initiators, providing strong evidence that these are active MGEs that replicate independently of their host. The sequential integration into the tRNA gene by distinct species of MGE and potential for mixing/synergy of their contents with genes deposited by previous insertion events is likely a beneficial aspect of this apparent method of acquiring defense systems.

Predicted active MGEs carrying this island, and others like it with different cargo, such as Dnd, BREX, and ferric citrate transport operons, all appear to be prophage-like, encoding prophage-like integrases, excisionases, and endonucleases of the HUH family involved in rolling circle replication initiation (Figure 5B, Supplementary Table S2). All these elements are flanked by 16-20 nt direct repeats (DRs), one located in the 3’ end of a tRNA gene, and the other at the presumed boundary of the element, likely, providing for excision by site-specific integrases/recombinases. These MGEs might mobilize upon prophage induction or phage infection and utilize (pro)phage proteins in *trans* to promote their own replication and cell-to-cell transmission via phage particles, interfering with (pro)phage reproduction while disseminating adaptive antiviral and resistance genes. Essentially, they could be acting as phage satellites akin to phage-induced chromosomal islands (PICIs) or Staphylococcal pathogenicity islands (SaPIs) (Quiles-Puchalt, Carpena et al. 2014, Fillol-Salom, Martínez-Rubio et al. 2018), but compared with those types of islands, the putative MGEs described encode a minimal number of recognizably phage-derived proteins, while delivering a large defensive payload.

Exploring this possibility, we subsequently recognized that, in addition to the defense systems, many of these MGEs carry phage-like “moron” modules, that is, operons encoding several small proteins, including RadC/ArdB antirestriction systems, CbtA/CbeA toxin-antitoxin systems, and several domains of unknown function (DUFs) found primarily in phages and located adjacent to one of the DRs at the opposite end of the element from the integrase (Figure 5A and 5B, Supplementary Table S2) (Katsiou, Nickel et al. 1999, Juhala, Ford et al. 2000, Heller, Tavag et al. 2017, Owen, Wenner et al. 2021, Kudryavtseva, Cséfalvay et al. 2023). Together, all these phage-like factors present in the predicted active defensive MGEs might promote their independent replication and interactions with host defenses and physiology that enable them to spread to other cells via helper phages. These observations support a general model of origin and evolution of defense islands, as successive waves of MGEs are inserted, immobilized, and maintained by selection for phage resistance (Figure 5B).

## Conclusions

Our analysis highlights the YprA helicase family as a prolific source of antiviral factors in the course of co-evolution between viruses and prokaryotic defenses, readily forming functional associations with a broad diversity of defense proteins, especially those shared with BREX and CoCoNuT systems. These helicases themselves adopt a remarkable diversity of domain architectures that likely contribute to the formation of multiprotein complexes involved in antivirus defense and, more generally, to the virus-host arms race. Even among the relatively well-characterized systems, such as DISARM and Druantia, considerable diversity remains to be studied, such as the Type IV CoCoNuTs. Several major branches, including the ARMADA, BRIGADE and TALON systems, were discovered in this work, but remain to be characterized in detail. Nevertheless, the experiments reported here demonstrated the powerful protective effect of ARMADA systems against a broad variety of phages. Furthermore, we show that YprA-centered defense systems are often encoded by predicted MGEs in which they form conserved defense gene arrays suggestive of coordinated activity. This hypothesis is supported by evidence of phage-specific synergy between the protective effects of ARMADA II and Druantia III. The study of the YprA-containing defense systems underscores the role of MGEs in spreading and likely facilitating the evolution of ever more complex defense systems in the perennial conflict with genetic parasites.

Many studies have effectively leveraged the tendency of prokaryotic immunity genes to aggregate into defense islands to reveal new classes of defense systems using a guilt-by-association approach. Here, we show how comprehensive phylogenetic analysis of abundant defensive proteins can also uncover large, previously unknown families of antiviral systems. We further present evidence of a widespread route of defense island origin and evolution whereby satellite phage-like MGEs, presumably packaged into helper phage particles, carry multiple defense systems, some of which synergize with each other, and integrate (often sequentially) into tRNA genes.

If this means of defense island mobilization is ultimately confirmed, it would be a striking demonstration of layered antiviral defense, in which these MGEs not only restrict phage genomes via the activity of specific defense systems, but also hijack their reproductive strategy, fitting within the framework of the ‘guns for hire’ concept of virus-immunity coevolution (Koonin, Makarova et al. 2020). In doing so, they could simultaneously reduce the yield of infectious phage particles while disseminating defense genes. In effect, these loci could transform the phage particle itself, one of the most formidable weapons in the viral arsenal, into a vehicle for antiviral protection.

## Methods

### Bioinformatics

The comprehensive search for YprA homologs was initiated by aligning with Muscle5 (Edgar 2022) sequences annotated as members of COG1205 (YprA) in a database of completely sequenced prokaryotic genomes maintained by our group. This alignment was then clustered, and each sub-alignment was used to produce a position-specific scoring matrix (PSSM). These PSSMs were then used as PSI-BLAST queries against the database as well as the non-redundant (nr) NCBI database to capture additional variants (E-value ≤⍰1e-5) (Altschul, Madden et al. 1997).

The YprA candidates identified during the search were clustered to a similarity threshold of 0.5 with MMseqs2 (Hauser, Steinegger et al. 2016), after which the representative sequences for each cluster were aligned with Muscle5. Approximately maximum-likelihood trees were built with the FastTree program (Price, Dehal et al. 2010) from these representative sequences. Detailed annotations of genome neighborhoods for the hits were generated by downloading their gene sequence, coordinates, and directional information from GenBank, as well as for 50 genes on each side of the hit. Domains in these genes were identified using PSI-BLAST against alignments of domains in the NCBI Conserved Domain Database (CDD) and Pfam (*E*-value 0.001). Comparing the results of the phylogenetic analysis with the genome neighborhood data distinguished the YprA-like hits from false positives. The clusters which were retained after this filtration step were re-clustered to similarity thresholds of 0.75 and finally 0.9, with alignment of cluster representatives, followed by phylogenetic and genome neighborhood analysis again after each step to filter out more false positive sequences. Although some eukaryotic YprA homologs were detected, these resembled the non-defensive YprA homologs in prokaryotes. Therefore, we excluded eukaryotic sequences from the final set of sequences used for our analysis to focus on the composition and contextual connections of prokaryotic YprA-containing systems (Supplementary Table S1). For the known and predicted defense systems and their genomic neighborhoods, we performed an extensive analysis of their domain composition using HHpred and often AlphaFold3, the latter of which produced, in most cases, predictions with accuracy comparable to experimentally solved structures (Gabler, Nam et al. 2020, Abramson, Adler et al. 2024).

### ARMADA, Druantia, and Zorya co-occurrence

Defense islands present in RefSeq v229 Bacteria and Archaea genomes were identified using PADLOC with a custom database including ARMADA system models (Payne, Todeschini et al. 2021). Only defense islands at least 10 ORFs away from the ends of contigs were included in the subsequent analyses. Redundant defense islands (both highly similar defense island gene content and island genetic context) were reduced by clustering the adjacent ORFs into groups based on their encoded protein sequence similarities using CD-HIT (with a 90% identity threshold over 80% pairwise alignment length), then grouping defense islands and their adjacent ORF families based on shared gene content, allowing up to three differences in gene content between families of unique genetic context and defense island composition.

### Bacteria and phages

*E. coli* strains, phages, and plasmids used in this study are listed in Supplementary Tables S3-5. *Escherichia coli* strain DH10B (New England Biolabs) was used for plasmid construction. Phage assays were performed in *E. coli* BL21-AI (Thermo Fisher Scientific) carrying plasmids expressing ARMADA, ARMADA mutant variants, YFP (negative control), or other defense systems. Cultures were grown in Lysogeny Broth (LB) at 37⍰°C with shaking at 180 rpm for liquid cultures, or on LB agar (LBA) plates for solid media. Where relevant, chloramphenicol was added to a final concentration of 25⍰µg/ml for plasmid maintenance.

### Cloning

ARMADA Type II was PCR-amplified from *E. coli* strain NCTC 12900 using Q5 High-Fidelity DNA Polymerase (New England Biolabs) and primers listed in Supplementary Table S4. PCR products were assembled into plasmid pBeloBAC11 using Gibson assembly (NEBuilder HiFi DNA Assembly Master Mix, New England Biolabs). Constructs were transformed into DH10B by electroporation and purified using the Monarch Plasmid Miniprep Kit (New England Biolabs). All plasmids were sequence-verified at Macrogen or Plasmidsaurus. Point mutations and gene deletions were introduced by Gibson assembly and confirmed by Sanger sequencing (Macrogen). Confirmed plasmids (Supplementary Table S3) were transformed into chemically competent BL21-AI cells using the Mix&Go! *E. coli* Transformation Kit (Zymo Research).

### Gene deletions in *E. coli*

The Δ*druEH* and Δ*armABCD* mutants of *E. coli* strain ATCC 8739 (NC_010468.1) were constructed using the λ Red recombineering method (Datsenko and Wanner 2000). Target operons were deleted from the first start codon to the last stop codon. A kanamycin resistance (*kanR*) cassette was amplified by PCR using Q5 High-Fidelity DNA Polymerase (NEB), with primers containing 50-to 60-bp homology arms corresponding to the flanking regions of the operons.

Electrocompetent cells were prepared from strains harboring the temperature-sensitive pKD46 plasmid. Cultures were grown in LB supplemented with ampicillin (100 µg/mL) at 30 °C. Expression of recombination machinery was induced by adding 0.2% arabinose at an OD_600_ between 0.3 and 0.4. Cultures were harvested at OD_600_ ≈ 0.6, washed with ice-cold 10% glycerol, and electroporated with 1,000 ng of purified PCR product (*kanR cassette*). After electroporation, cells were recovered in SOC at 37 °C for 4 h to allow recombination and pKD46 plasmid curing. Cells were then plated on kanamycin (50 µg/mL) plates and incubated overnight at 37 °C. Loss of pKD46 was confirmed based on ampicillin sensitivity.

To remove the kanR cassette, verified mutant strains were transformed with the FLP recombinase-expressing plasmid pCP20. Electroporated cells were recovered in SOC at 30⍰°C for 1 h, then plated on LB agar with ampicillin (100 µg/mL) and incubated overnight at 30⍰°C. Single colonies were restreaked onto antibiotic-free LB agar and incubated at 40⍰°C overnight to induce FLP recombinase activity and eliminate the pCP20 plasmid. Clones were tested for antibiotic sensitivity to both kanamycin and ampicillin and verified by colony PCR and Sanger sequencing.

### Efficiency of plating

To determine the efficiency of plating (EOP) of different phages as part of Figure 2, overnight cultures of *E. coli* BL21-AI containing the plasmid encoding ARMADA, ARMADA mutants, or YFP were mixed with 0.6% (w/v) LB soft agar and overlaid on LBA plates. Ten-fold serial dilutions of phage stocks were spotted onto the lawns and plates incubated overnight at 37⍰°C. Phage plaques were counted and used to calculate the EOP relative to the YFP-expressing control.

To determine the EOP of *E. coli* NC_010468.1 in Figure 4, cells were challenged with phages from the Basel phage collection (Maffei, Shaidullina et al. 2021) and additional isolates using a soft agar overlay assay. As part of the assay, overnight cultures were diluted 1:100 in LB and cultured at 37⍰°C with shaking until reaching OD_600_ ≈ 0.6. Cells were harvested by centrifugation (3,000⍰× ⍰g for 5⍰min at 4⍰°C), resuspended in 200⍰µL of cold LB,and maintained on ice. For soft agar overlays, 100⍰µL of bacterial suspension was mixed with 10⍰mL of soft agar (0.35%, pre-warmed to 55°C) and poured onto pre-warmed LB agar plates. Plates were gently swirled to evenly distribute the soft agar and left to solidify at room temperature for up to 30 min.

Phage dilutions were prepared in LB, and 3⍰µL of each dilution were spotted onto the surface of the soft agar layer. Plates were incubated at 37⍰°C overnight. Plaques were counted the next day. For lytic phages that did not produce distinct plaques, the lowest dilution showing a clear lysis halo was used for quantification. Plaque size was estimated by measuring the radius of three individual plaques per condition.

### Liquid infection assays

To assess phage infection in liquid culture in Figure 2, overnight cultures of E. coli BL21-AI containing the plasmid encoding ARMADA or YFP were diluted to OD_600_⍰= ⍰0.1 in LB with chloramphenicol and distributed into wells of a 96-well flat-bottom plate. Phages were added at different multiplicity of infection (MOI), or LB alone for mock controls. Plates were incubated at 37°C with shaking at 200 rpm in a CLARIOstar Plus plate reader (BMG Labtech), with OD_600_ measured every 10 minutes for 500 minutes.

To assess phage infection in liquid culture in Figure 4, wild-type and mutant strains of E. coli NC_010468.1 were subjected to liquid culture phage challenge at different multiplicities of infection (MOIs). Overnight cultures were diluted 1:100 in fresh LB and grown at 37⍰°C until OD_600_ ≈ 0.3. For each condition, 180⍰µL of bacterial culture was mixed with 20⍰µL of phage lysate, previously diluted to achieve final MOIs of 0.1, 1, or 10. Samples were transferred to a 96-well plate and incubated in a BioTek Synergy H1 microplate reader at 37°C. The OD_600_ was recorded every 10 min for 9 h to monitor bacterial growth and phage-induced lysis.

### Membrane permeability assay with propidium iodide

Overnight cultures of BL21-AI containing the plasmid encoding ARMADA or YFP were diluted 1:100 in LB with chloramphenicol and grown at 37⍰°C with shaking until OD_600_⍰= ⍰0.4. Cultures were then diluted to OD_600_⍰= ⍰0.05 in LB with chloramphenicol and 1% (v/v) DMSO. Propidium iodide (PI) (Sigma-Aldrich) was added to a final concentration of 5⍰µg/ml and cultures were incubated for 1 hour before phage addition. Phages were added at an MOI of 0.01 or 10. OD_600_ and PI fluorescence (excitation: 544⍰nm; emission: 612⍰nm) were measured every 10 min for 500 minutes using a CLARIOstar Plus plate reader. PI fluorescence was normalised to OD_600_.

### Metabolic activity assay with resazurin

Cultures of BL21-AI containing the plasmid encoding ARMADA or YFP were prepared as for the membrane permeability assay. Resazurin (Sigma-Aldrich) was added to a final concentration of 3µg/ml after 1 hour incubation at 37⍰°C with shaking and phages were added at MOI 0.01 or 10. OD_600_ and resazurin fluorescence (excitation: 560⍰nm; emission: 590⍰nm) were recorded every 10 minutes for 500 minutes. Resazurin signal was normalised to OD_600_.

### Statistical analysis

Unless stated otherwise, the presented results are the mean of three biological replicates. For the data shown in Figure 2, statistical analyses were performed using GraphPad Prism v9.2.0, and values are presented as mean ± standard deviation. One-way ANOVA with multiple comparison correction was used to access statistical significance, with p-value <0.05 considered significant. Synergy analyses were conducted in R version 4.4.2. One-sided unpaired t-tests were used to assess statistical significance following confirmation of data normality.

EOP and SFC values were calculated relative to the double ARMADA Type II/Druantia Type III mutant. Statistical significance was assessed using an unpaired t-test with the alternative hypothesis ‘greater’ in R version 4.4.2. Epistatic coefficients, representing the interaction strength between defense systems, were calculated as |log10 (EOP{system1+system2})| – |log10(EOP{system1})| – |log10(EOP{system2})| as described previously (Wu, Garushyants et al. 2024). The coefficient was set to 0 if the mean EOP of WT strains for a given virus was ≥ 1×10^-2^.

To evaluate the impact of interactions between defense systems in the liquid assay, we calculated the area under the curve (AUC) for strains carrying individual systems and their combinations. For the AUC calculation, the optical density at the start of the experiment was subtracted from each data point. If the AUC of a system combination exceeded the sum of the individual AUCs (expected additive value), the systems were considered to have a synergistic protective effect.

---

**Supplementary Figure S1.**
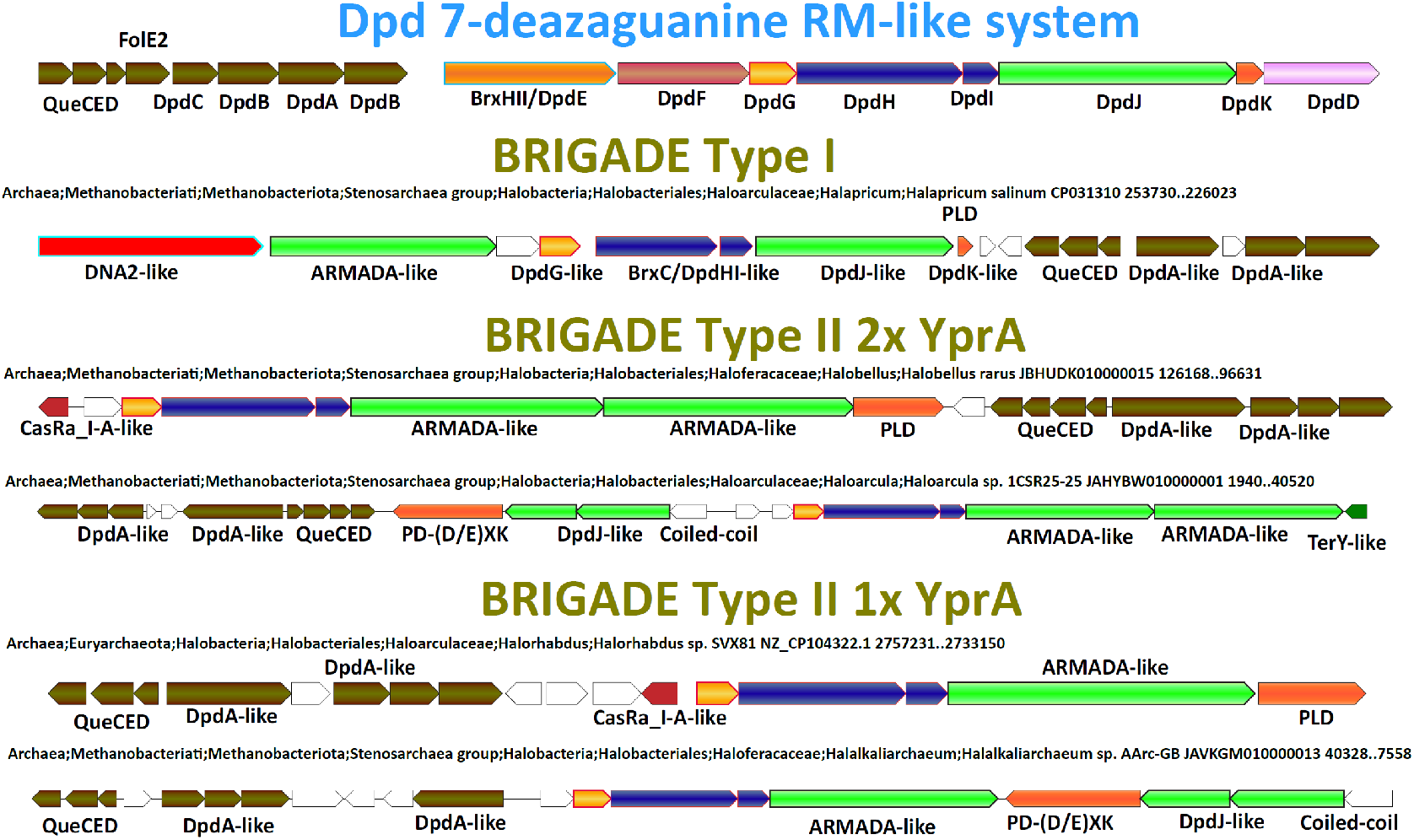
Comparison of Dpd with BRIGADE systems found in halobacteria. Canonical Dpd systems encompass a 7-deazaguanine derivative biosynthesis and insertion operon, as well as BREX-like factors DpdE/BrxHII and DpdHI/BrxC (BrxHII is shared with DISARM and ARMADA), a YprA family helicase (DpdJ), and additional factors involved in restriction of non-hypermodified DNA. BRIGADE systems share many of these factors, including many, but not all of the genes in the 7-deazaguanine derivative biosynthetic cluster, DpdG, DpdHI/BrxC, at least one YprA homolog, and PLD superfamily endonucleases. BRIGADE Type II also usually includes a CasRa-like regulatory protein and has an alternative nuclease in an operon with a DpdJ-like YprA homolog when the PLD nuclease is not present. In contrast, BRIGADE Type I is almost always associated with a small PLD nuclease and often with a large DNA2-like helicase/nuclease fusion protein.

**Supplementary Figure S2.**
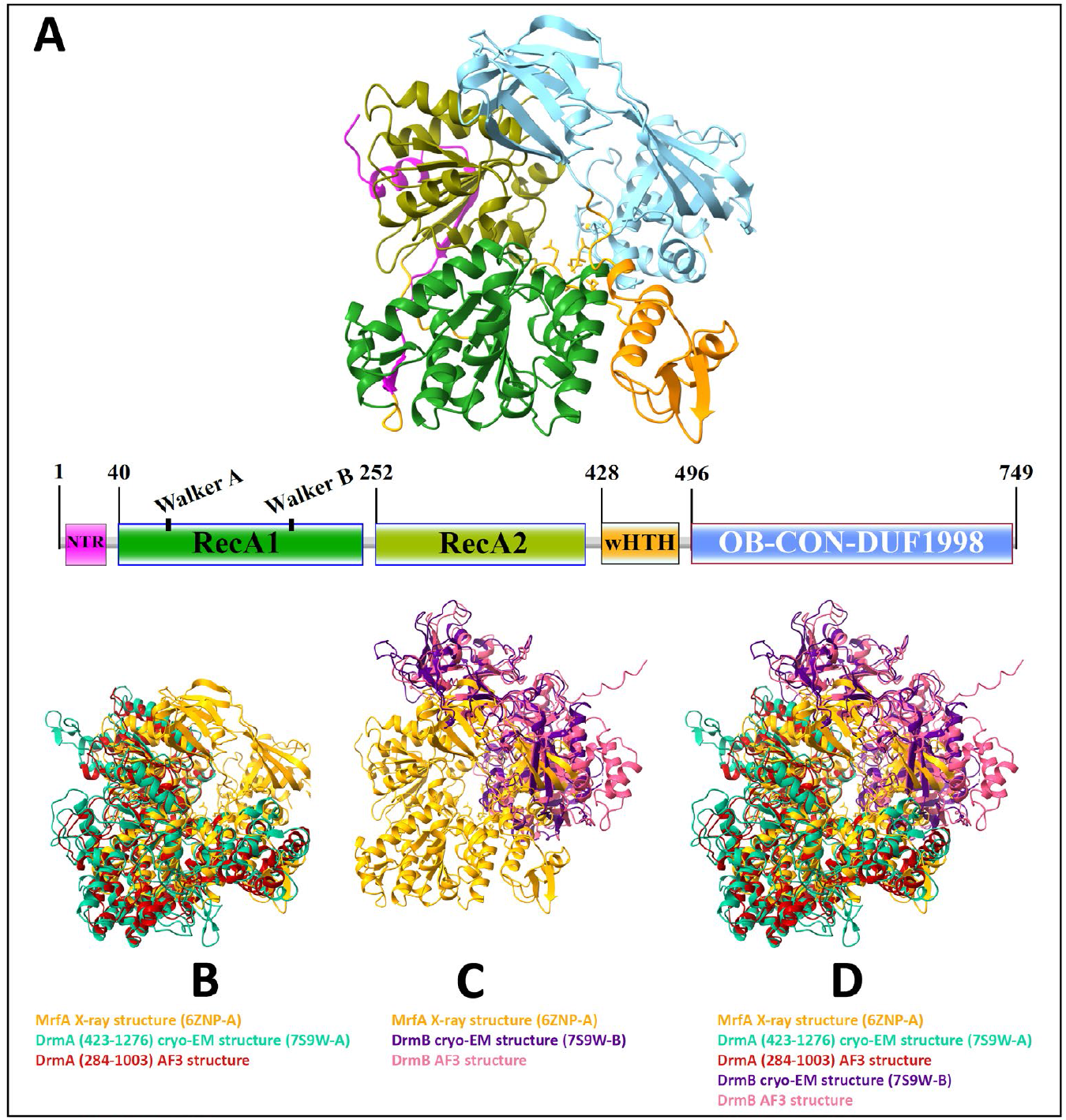
Experimental and AF3 predicted structures of MrfA and DrmAB. (A) MrfA X-ray crystallographic structure and representation of domain architecture. The domains annotated are the N-terminal region (NTR, magenta), the first RecA-like domain containing the Walker A/B motifs (RecA1, dark green), the second RecA-like domain (RecA2, olive), a wHTH domain (orange), and the canopy-forming OB-CON-DUF1998 domain array (blue). (B) Superpositions of MrfA (gold), the DrmA helicase structure solved in complex with DrmB (teal), DrmA only shown here with N and C-terminal truncations for clarity, and a representative DrmA AF3 predicted structure (red), with similar N and C-terminal truncations. (C) Superpositions of MrfA (gold), the DrmB OB-CON-DUF1998 structure solved in complex with DrmA by Bravo et al. (purple), DrmB only shown here for clarity, and a representative DrmB AF3 predicted structure (pink). (D) Superpositions of MrfA (gold), the DrmAB helicase structure solved by Bravo et al. (DrmA: teal, DrmB: purple), with N and C-terminal truncations of DrmA for clarity, a representative DrmA AF3 predicted structure (red), with similar N and C-terminal truncations, and a representative DrmB AF3 predicted structure (pink).

**Supplementary Figure S3.**
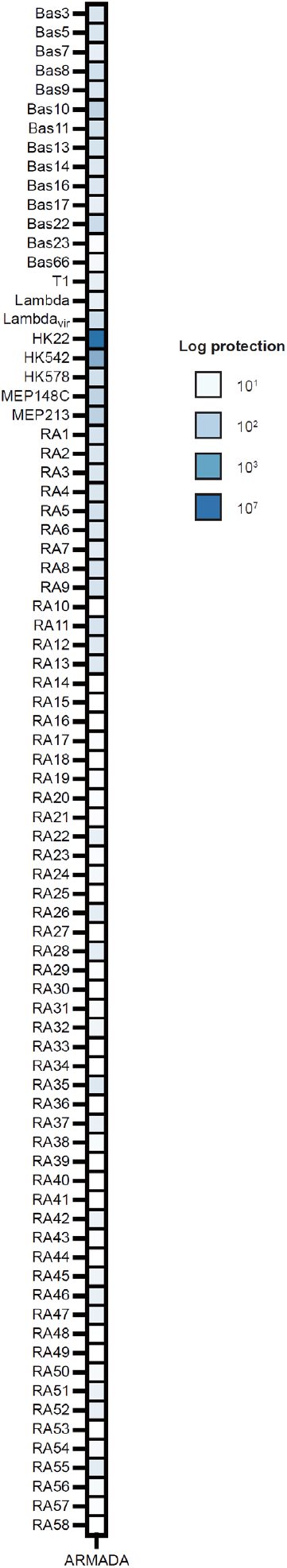
Complete efficiency of plating (EOP) results for the ARMADA anti-phage protection experiments shown in Figure 2. Log-protection (reduction in phage titer) conferred by ARMADA, expressed from plasmid pBeloBAC11 in BL21-AI, against a panel of 80 phages

